# Analysis of brain atrophy and local gene expression in genetic frontotemporal dementia

**DOI:** 10.1101/2019.12.11.872143

**Authors:** Andre Altmann, David M Cash, Martina Bocchetta, Carolin Heller, Regina Reynolds, Katrina Moore, Rhian S Convery, David L Thomas, John C van Swieten, Fermin Moreno, Raquel Sanchez-Valle, Barbara Borroni, Robert Laforce, Mario Masellis, Maria Carmela Tartaglia, Caroline Graff, Daniela Galimberti, James B Rowe, Elizabeth Finger, Matthis Synofzik, Rik Vandenberghe, Alexandre de Mendonça, Fabrizio Tagliavini, Isabel Santana, Simon Ducharme, Chris R Butler, Alex Gerhard, Johannes Levin, Adrian Danek, Giovanni Frisoni, Roberta Ghidoni, Sandro Sorbi, Markus Otto, Mina Ryten, Jonathan D Rohrer, on behalf of the Genetic FTD Initiative, GENFI

## Abstract

Frontotemporal dementia (FTD) is a heterogeneous neurodegenerative disorder characterized by neuronal loss in the frontal and temporal lobes. Despite progress in understanding which genes are associated with the aetiology of FTD, the biological basis of how mutations in these genes lead to cell loss in specific cortical regions remains unclear. In this work we combined gene expression data for 16,772 genes from the Allen Institute for Brain Science atlas with brain maps of gray matter atrophy in symptomatic *C9orf72*, *GRN* and *MAPT* mutation carriers obtained from the Genetic FTD Initiative study. No significant association was seen between (*C9orf2*, *GRN* and *MAPT* expression and the atrophy patterns in the respective genetic groups. Between 1,000 and 5,000 genes showed a negative or positive correlation with the atrophy pattern within each individual genetic group, with the most significantly associated genes being *TREM2*, *SSBP3* and *GPR158* (negative association in *C9orf72*, *GRN* and *MAPT* respectively) and *RELN, MXRA8* and *LPA* (positive association in *C9orf72*, *GRN* and *MAPT* respectively). An overrepresentation analysis identified a negative correlation with genes involved in mitochondrial function, and a positive correlation with genes involved in vascular and glial cell function in each of the genetic groups. After adjusting for spatial autocorrelation, a set of 423 and 700 genes showed significant positive and negative correlation, respectively, with atrophy patterns in all three maps. The gene set with increased expression in spared cortical regions was enriched for neuronal and microglial genes, while the gene set with increased expression in atrophied regions was enriched for astrocyte and endothelial cell genes. Our analysis suggests that these cell types may play a more active role in the onset of neurodegeneration in FTD than previously assumed, and in the case of the positively-associated cell marker genes, potentially through emergence of neurotoxic astrocytes and alteration in the blood-brain barrier respectively.

**Abbreviated summary:** Altmann et al. investigated the concordance between spatial cortical gene expression in healthy subjects and atrophy patterns in genetic frontotemporal dementia. They found that elevated gene expression of endothelial cell and astrocyte-related genes in regions with atrophy, suggesting a role of these cell types in the aetiology of frontotemporal dementia.

## Introduction

Frontotemporal dementia (FTD) is a heterogeneous neurodegenerative disorder characterized by neuronal loss in the frontal and temporal lobes, with clinical symptoms including behavioural, language and motor deficits (Seelaar *et al*., 2011). Around 30% of FTD is familial, most commonly caused by autosomal dominant genetic mutations in one of three genes: progranulin (*GRN*), microtubule-associated protein tau (*MAPT*) or chromosome 9 open reading frame 72 (*C9orf72*) (Rohrer *et al*., 2009). Despite progress in understanding the pathophysiological basis of genetic FTD, the biological basis of how mutations in these genes leads to cell loss in specific cortical regions and subsequently to specific clinical phenotypes is unclear.

An alternative approach to elucidating the molecular biology of autosomal dominant FTD is to study the gene expression profiles of brain regions which are atrophic in symptomatic mutation carriers. This approach has been enabled by publicly available data from the Allen Institute for Brain Science which features post-mortem high-resolution brain-wide gene expression data (i.e., the Allen Atlas) from cognitively normal individuals (Hawrylycz *et al*., 2012). In recent years the Allen Atlas has been successfully integrated with brain maps obtained from case-control studies. For instance, in the case of neurodegenerative disorders, one study investigated the link between gene expression and both regional patterns of atrophy and amyloid deposition, finding a positive correlation of *APP* gene expression and amyloid (Grothe *et al*., 2018), whilst another study showed that expression of the *MAPT* gene was associated with changes in functional connectivity in Parkinson’s disease (Rittman *et al*., 2016).

In this work we combine gene expression data from the Allen Atlas with brain maps of gray matter atrophy in symptomatic *C9orf72, GRN* and *MAPT* mutation carriers from the Genetic FTD Initiative (GENFI) study compared with non-carriers (Cash *et al*., 2017). The aim of this study was to investigate the molecular basis of the atrophy pattern in mutation carriers. We firstly investigated the spatial overlap between gray matter atrophy in each of the three genetic groups and the gene expression of the corresponding gene. We then aimed to identify which genes showed a high spatial correspondence between their expression throughout the brain and the atrophy pattern in each genetic group. We hypothesized that these genes or groups of genes may implicate molecular processes or brain cell types that explain why these regions are particularly vulnerable in FTD.

## Methods

### Allen Human Brain Atlas Data

We used the brain-wide microarray gene expression data generated by the Allen Institute for Brain Science (downloaded from http://human.brain-map.org/) (Hawrylycz *et al*., 2012). The dataset consists of a total of 3,702 microarray samples from six donors (one female). Each sample comprised 58,692 gene probes and provides coordinates in MNI152 space. The gene expression data have been normalized and corrected for batch effects by the Allen Institute (‘TECHNICAL WHITE PAPER: MICROARRAY DATA NORMALIZATION’, 2013). We first restricted the set of samples to the left hemisphere (six in total) and to cortical regions based on the provided slab type (‘*cortex*’), retaining only samples with a maximal distance of 3 mm to a cortical region of interest (ROI) obtained by a parcellation (Cardoso *et al*., 2015) of the study template used in Cash *et al*. (2017). After additionally removing samples being annotated by the Allen Institute as non-cortical samples (e.g., CA1 field), 1,248 microarray samples were available in total. Next, as previously described (Richiardi *et al*., 2015) we reannotated all microarray probe sequences with gene names using Re-Annotator (Arloth *et al*., 2015). We excluded probes that sampled more than one gene (N=1,512), were mapped to intergenic regions (N=5,013) or could not be mapped to any genomic region (N=1,569), leaving 50,598 probes covering 19,980 unique genes. Furthermore, we removed probes that were marked as expressed in less than 300 of the 1,248 cortical samples (N=13,941). Thus, the analysis was carried out using 36,657 microarray probes covering 16,772 distinct genes.

### Image data preparation

In order to quantify the amount of atrophy in carriers of FTD mutations, we used results from a voxel-based morphometry (VBM) analysis of the GENFI dataset (Cash *et al*., 2017). In particular, in this analysis we used the maps showing the voxel-wise t-statistic (t-maps) comparing symptomatic mutation carriers (*MAPT*: N=10; *GRN*: N=12; *C9orf72*: N=25) to non-carriers (N=144) (Figure 1; top). Here, higher t-scores signify higher average atrophy in the symptomatic group analysis. A mean bias corrected image from all the normalized T1 images in the GENFI study served as a study template. This template was warped into MNI space using the non-rigid registration based on fast free form deformation implemented in `NiftyReg` (version 10.4.15) (Modat *et al*., 2010). The obtained transformation was then applied to each of the three t-maps. For each MNI coordinate of the eligible cortical gene expression samples we located the corresponding voxel in the t-map and obtained the t-value centered on those MNI coordinates. This procedure was carried out for each of the three t-maps and resulted in a 1,248 by three matrix, i.e., each gene expression sample was linked to three t-scores from the VBM analysis (one for each FTD gene).

**Figure 1:**
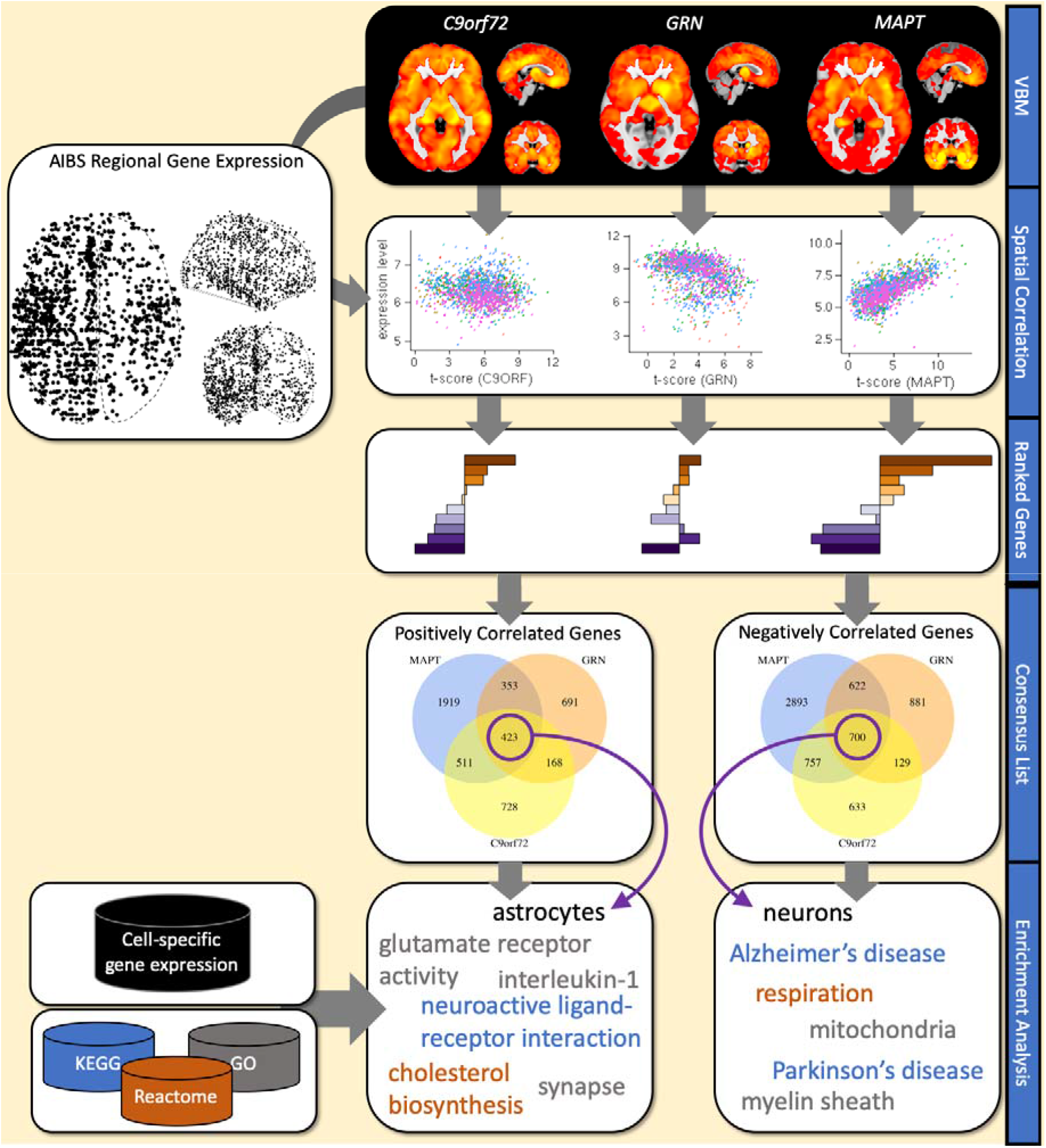
Analysis overview. Starting point of the analysis are the statistical maps from voxel-based morphometry analyses (VBM) comparing healthy controls and symptomatic FTD mutation carriers in *C9orf72, GRN* and *MAPT* from *Cash et al*. (2017). For each statistical map the spatial correlation with the expression levels 16,772 genes (represented by 36,657 gene expression probes) is computed using data from the Allen Institute for Brain Science (AIBS) gene expression atlas. The resulting gene ranking provides lists of genes that are either significantly (P_FDR_<0.05) positively or negatively correlated with the atrophy pattern. Two consensus lists were generated from the three lists of positively correlated and negatively correlated genes, respectively. Resulting gene lists were analyzed for enrichment of signature genes for brain cell types, such as neurons, microglia or astrocytes, and biological pathways.

### Association analysis

The overall analysis is depicted in Figure 1. We analyzed the association between atrophy and gene expression in a quasi-non-parametric fashion. For a given atrophy map and a given probe, we approximated the Spearman (or rank) correlation (ρ) between the local t-score and the gene expression level separately for each of the six donors while accounting for spatial autocorrelation. This was achieved by first converting t-scores and expression levels into ranks, respectively, and using linear regression to approximate the Spearman rank correlation (Conover and Iman, 1981); and second by adjusting this linear model using spatial eigenvectors that account for the spatial relationship between samples. More precisely, we used the method of spatial eigenvector mapping (reviewed in (Dormann *et al*., 2007)) in order to adjust for spatial autocorrelation. In this approach, one first checks the residuals of the linear model for presence of spatial autocorrelation using Moran’s I (Moran, 1950). If there was evidence for spatial autocorrelation, then the first spatial eigenvector was added as a confound variable to the linear model. Next, if the residuals of the resulting extended linear model continued to show evidence for spatial autocorrelation, then further eigenvectors were added until the residual was free from evidence of spatial autocorrelation. To accelerate the process, spatial eigenvectors were added to the model in batches of five and up to 150 spatial eigenvectors were considered. Spatial eigenvectors for this analysis were obtained from the ten-nearest neighbor graph constructed using the geodesic distances between gene expression samples, which were generated for each donor brain. From the resulting linear model, we extracted the t-scores and the corresponding p-value for the association between atrophy and gene expression. Next, we combined the six p-values into a single meta p-value using the weighted sum of Z scores (Stouffer’s) method. On purpose, we restricted weights to −1 and +1 to only indicate the direction of association, and to avoid over-emphasizing the impact of donors with more gene expression samples. This procedure was carried out for each for the 36,657 probes and each of the three genetic group atrophy maps. P-values in each of the three resulting lists were corrected for multiple testing using the method by Benjamini and Hochberg (1995) for FDR.

Significantly positively correlated genes (i.e., higher gene expression is linked to higher atrophy) were those where any probe targeting the gene reached an FDR-corrected p-value < 0.05 and a positive z-score; likewise significantly negatively correlated genes (i.e., higher gene expression is linked to lower atrophy) were required to have an FDR-corrected p-value < 0.05 and a negative z-score for any of the probes targeting that gene. We also created two overlap lists, one containing the overlap of genes in the three positive lists, the other the overlap of genes in all three negative lists. In the following we refer to these two lists as *consensus* lists.

### Overrepresentation analysis

In order to identify cellular pathways or cellular processes and cell type signature genes that may be enriched in the significant gene lists, we conducted an overrepresentation analysis. We obtained the following gene sets from the MSigDB database version 6.1 (date accessed 11/23/2017): i) Gene Ontology (N=5,917 sets), ii) REACTOME (N=674 pathways) and iii) KEGG (N=186 pathways). We used Fisher’s exact test to compute the p-value for overrepresentation of genes in a given set. All tests were carried out using the 16,772 cortex expressed genes as the background set. For each of the three gene lists (i.e., one per FTD gene) and the two consensus lists we corrected the p-values using the FDR correction based on Benjamini and Hochberg across all 6,777 gene sets.

In order to determine whether the expressed genes implicate a specific class of brain cell types, in addition to enrichment analysis for GO terms and pathways, we conducted an enrichment analysis for marker gene lists for six brain cell-types based on RNA sequencing of purified human cells (Zhang *et al*., 2016). In brief, we generated cell-type specific lists from the available average expression levels per cell-type: only genes with expression levels exceeding 2.5 fragments per kilobase (FPKM) were included and genes were required to show an enrichment of at least 3.0 (i.e., FPKM in target cell-type divided by average FPKM in non-target cell-types).

### Expression Weighted Cell-Type Enrichment analysis

In an additional analysis we sought to identify potential brain cell types that were implicated by all three FTD genes. To this end we conducted Expression Weighted Cell-type Enrichment (EWCE) analysis (Skene and Grant, 2016) on the two consensus lists using a recently published dataset of brain single-cell sequencing data in the mouse brain that identified 265 different cell types (www.mousebrain.org) (Zeisel *et al*., 2018). From this dataset we removed 76 cell types that were not directly brain related (e.g., cell belonging to enteric nervous system or the spinal cord), leaving 189 different cell-type signatures. Each of the cell types is also attributed with a high-level annotation (astrocytes, ependymal, immune, neurons, oligos, vascular). In brief, from the single cell mouse dataset we used only genes that had a unique human homolog (1-to-1 mapping). Then, we analyzed the two consensus lists separately for high-level cell type enrichment using EWCE with correction for gene length and GC content. EWCE was executed twice, once using high-level annotations and once using the 189 cell-type annotations. P-values are based on 100,000 permutations and enrichment P-values were corrected for multiple testing using the Benjamini and Hochberg method for FDR. We used the available R package for EWCE.

### Interpretation of association, overrepresentation and EWCE analyses

Overall, the type of gene expression profiling used in Allen Atlas is based on measuring bulk expression of tissue samples, i.e., a group of diverse cell types is sampled at once and the resulting expression profile represents the group average of this set of cells and their states. Thus, in these analyses, genes positively correlated with atrophy indicate potential cellular processes and cell types that promote atrophy in genetic FTD. Conversely, genes that are negatively correlated with atrophy indicate potential cellular processes and cell types that confer resilience to disease-related neurodegeneration.

### Data availability

Statistical maps of the VBM analysis were obtained from Cash *et al*. (2017) and are available upon request from the lead author of that study. Cortical gene expression data were obtained from the Allen Institute of Brain Sciences and can be obtained at http://human.brain-map.org/. Code for pre-processing expression data and conducting the regional correlation analysis with adjustment for spatial autocorrection can be accessed through: https://github.com/andrealtmann/AIBS_FTD.

## Results

We tested 16,776 genes (from 36,657 microarray probes) for their association with atrophy pattern across the cortex in genetic FTD (Figure 1). The numbers of significant probes (P_FDR_<0.05) and genes with their direction for each of the three FTD genes are listed in Table 1. The association results per probe are available as supplementary material (Dataset S1).

**Table 1:**
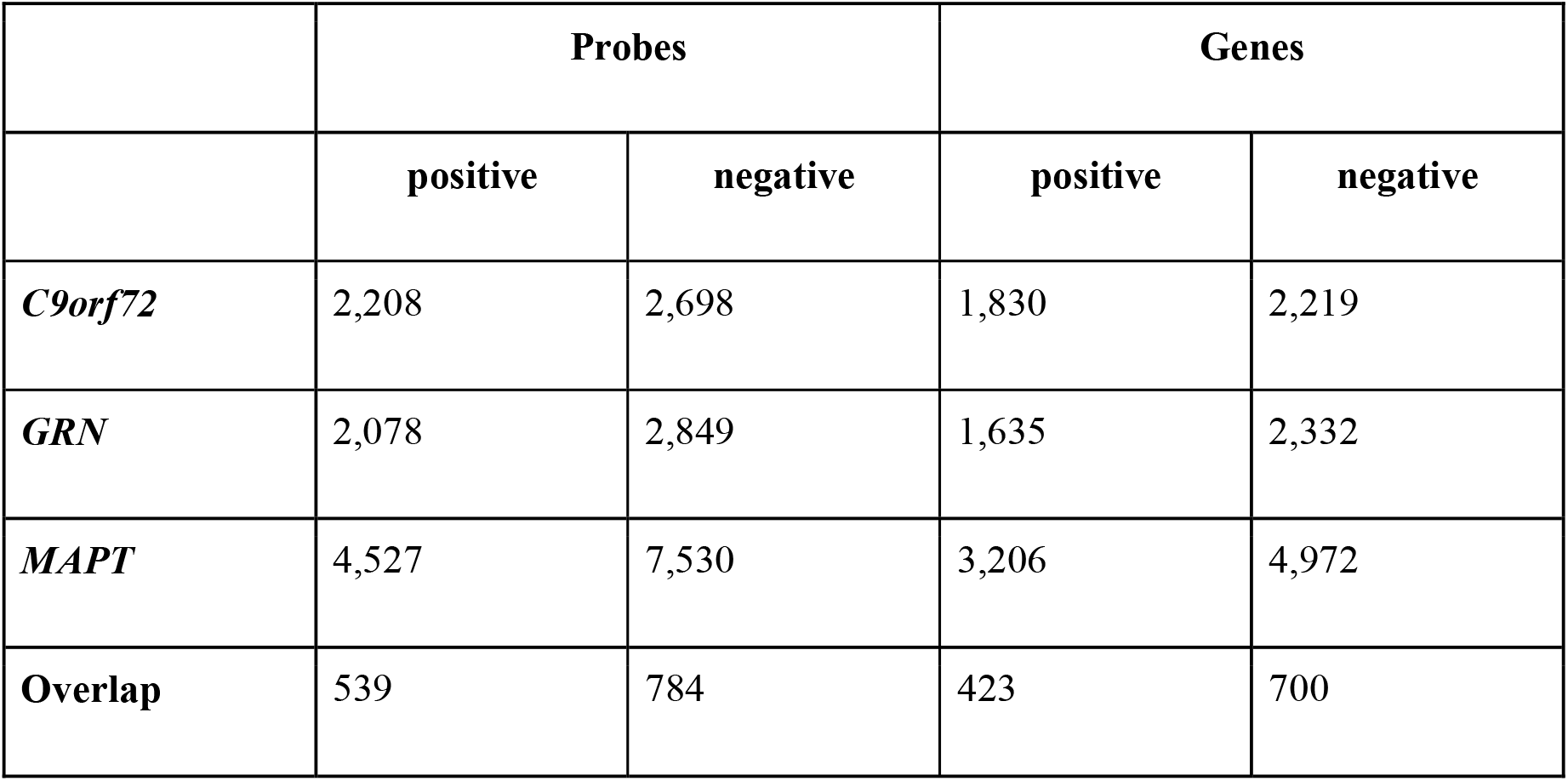
Numbers of significant probes and genes and their direction of association after Benjamini and Hochberg correction for FDR. Rows correspond to the three genes and an overlap between all three lists.

|  | Probes |  | Genes |  |
| --- | --- | --- | --- | --- |
|  | positive | negative | positive | negative |
| *C9orf72* | 2,208 | 2,698 | 1,830 | 2,219 |
| *GRN* | 2,078 | 2,849 | 1,635 | 2,332 |
| *MAPT* | 4,527 | 7,530 | 3,206 | 4,972 |
| **Overlap** | 539 | 784 | 423 | 700 |

### C9orf72

The strongest association between *C9orf72* expression and the atrophy pattern in symptomatic C9orf72 repeat extension carriers was measured with microarray probe CUST_7641_PI416261804, which showed a non-significant negative association z=-0.27 (P_FDR_=0.23; Figure 2).

**Figure 2:**
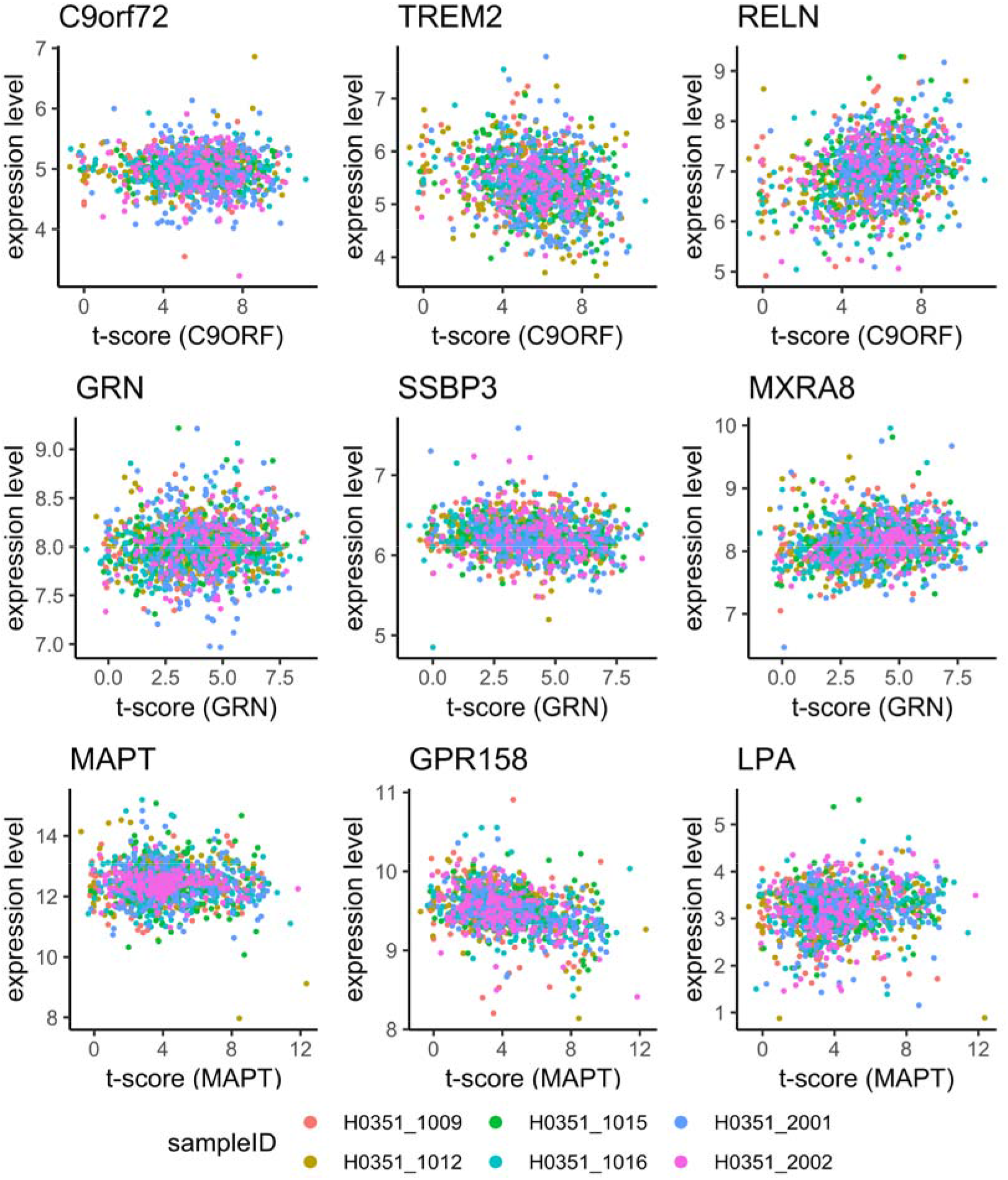
Selected scatterplots between t-scores and gene expression levels. Rows correspond to the three different FTD genes (*C9orf72, GRN* and *MAPT*). X-axes show to the t-value in the corresponding statistical VBM maps (Figure 1) from Cash *et al*. (2017) at positions where microarray samples were obtained in the Allen Atlas; the y-axes represent the expression levels in the Allen Atlas after adjustment for spatial autocorrelation for the gene named at the top of the scatter plot. Each point represents one microarray sample and the color indicates the donor ID. In the first column expression of the corresponding FTD genes is studied: *C9orf72* (CUST_7641_PI416261804), *GRN* (CUST_13046_PI416261804) and *MAPT* (A_24_P224488). The second column shows the most strongly negatively correlated genes, i.e., genes with high expression in brain regions showing little atrophy: *TREM2* (A_23_P167941), *SSBP3* (A_23_P500333) and *GPR158* (A_24_P349117). The third column shows the genes where expression level and atrophy positively correlate the strongest: *RELN* (A_24_P309095), *MXRA8* (A_23_P32444), and *LPA* (A_23_P95221).

The most negatively associated gene was *TREM2* (Triggering Receptor Expressed On Myeloid Cells 2; represented by probe A_23_P167941; z=-6.26; P_FDR_=6.94e-07; Figure 2; Table 2; Dataset S1). Top-ranked gene sets based on the significantly negatively correlated genes include genes related to mitochondria (GO_MITOCHONDRIAL_PART; OR=3.00; P_FDR_=1.77e-35) and the respiratory chain (GO_RESPIRATORY_CHAIN; OR=9.92; P_FDR_=5.91e-19; Dataset S2). Notably, KEGG pathways for neurodegenerative disorders were highly enriched (KEGG_PARKINSONS_DISEASE OR=7.35 P_FDR_=8.74e-20; KEGG_HUNTINGTONS_DISEASE OR=4.99 P_FDR_=8.18e-18; KEGG_ALZHEIMERS_DISEASE OR=4.15 P_FDR_=4.38e-14). Among brain cell types, there was a strong enrichment for neuronal genes (OR=1.7; P_FDR_=1.13e-12) and microglia (OR=1.59; P_FDR_=2.53e-06; Dataset S3).

**Table 2:**
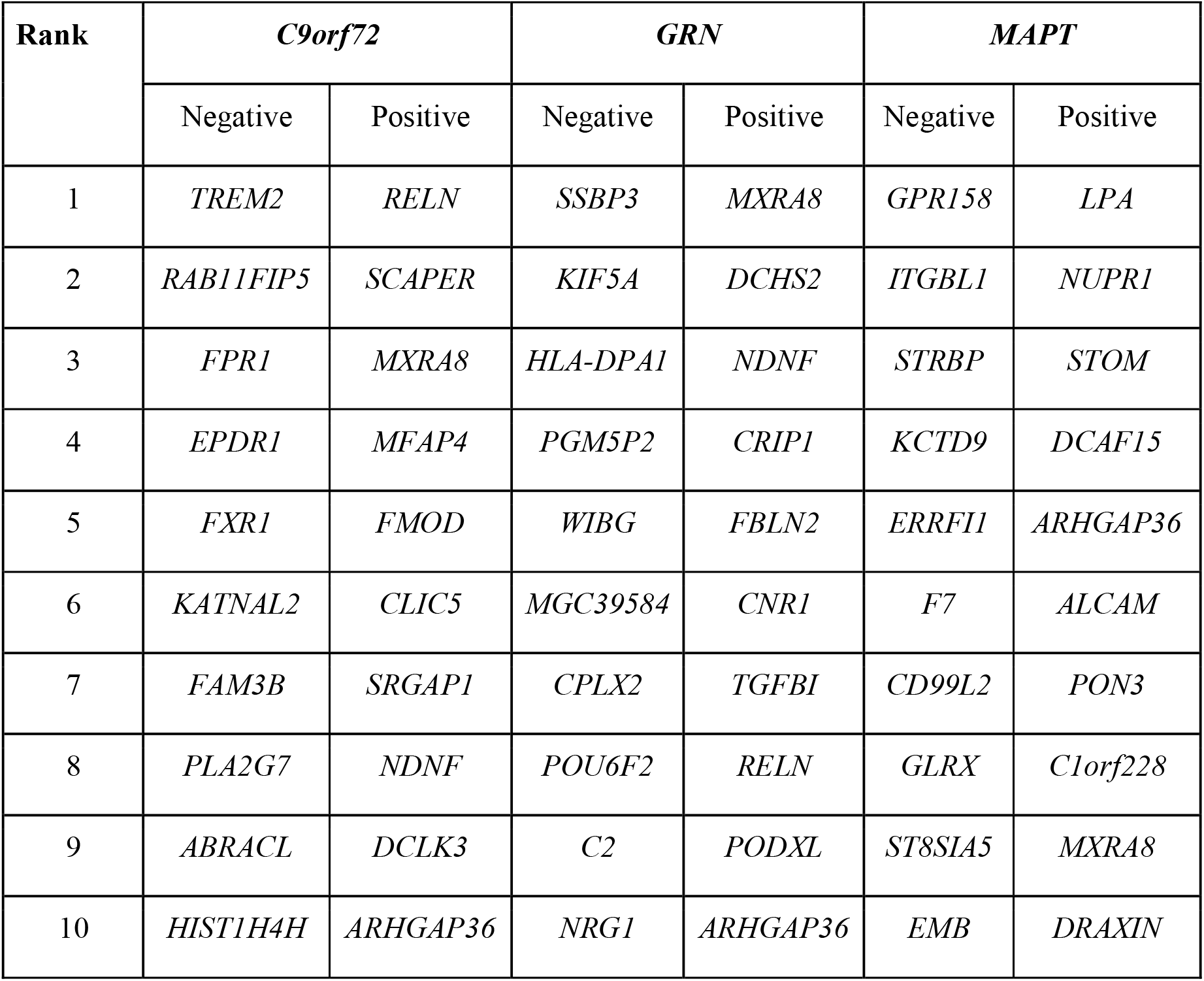
Top ten most positively and negatively correlated genes with atrophy patterns in genetic FTD.

The most significantly positively correlated gene was *RELN* (reelin; represented by probe A_24_P309095; z=7.94; P_FDR_=1.92e-11; Figure 2, Table 2). Top-ranked GO terms for positively correlated genes included vascular development and glial cell differentiation (GO_REGULATION_OF_VASCULATURE_DEVELOPMENT OR=2.28 P_FDR_=9.95e-04; GO_GLIAL_CELL_DIFFERENTIATION OR=2.66 P_FDR_=1.13e-03; Dataset S2). Among brain cell types, there was a strong enrichment for genes associated with oligodendrocytes (OR=9.37; P_FDR_=3.1e-54), endothelial cells (OR=2.4; P_FDR_=3.63e-10) and mature astrocytes (OR=1.62; P_FDR_=1.23e-05; Dataset S3).

### GRN

The strongest association between *GRN* expression and the atrophy pattern in symptomatic GRN mutation carriers was measured with microarray probe CUST_13046_PI416261804, which showed a non-significant positive association z=1.31 (P_FDR_=0.13; Figure 2).

The most significantly negatively correlated gene was *SSBP3* (Single Stranded DNA Binding Protein 3; Figure 2; Table 2). For the negatively correlated genes, significantly enriched gene sets featured the respiratory chain (e.g., REACTOME’s RESPIRATORY_ELECTRON_TRANSPORT OR=5.04 P_FDR_=6.13e-06; Dataset S2). As with C9orf72, the three KEGG pathways for neurodegenerative disorders were enriched (Alzheimer’s OR=2.38 P_FDR_=3.64e-03; Parkinson’s OR=3.09 P_FDR_=2.17e-04; Huntington’s OR=2.26 P_FDR_=5.32e-03). Among brain cell types, there was a strong enrichment for neuronal genes (OR=1.71; P_FDR_=2.06e-13) and microglia (OR=1.68; P_FDR_=4.10e-08; Dataset S3).

The most significantly positively associated gene was *MXRA8* (Matrix Remodeling Associated 8; Figure 2; Table 2). Top-ranked GO terms (Dataset S2) for positively correlated genes are related to the extracellular matrix (GO_EXTRACELLULAR_MATRIX OR=3.79 P_FDR_=1.25e-18), vascular development, (GO_VASCULATURE_DEVELOPMENT OR=2.68 P_FDR_=4.16e-11), and response to wounding (GO_RESPONSE_TO_WOUNDING OR=2.45 P_FDR_=9.19e-10). Again, genes related to oligodendrocytes showed the strongest enrichment (OR=9.47; P_FDR_=6.1e-53), followed by mature astrocytes (OR=3.19; P_FDR_=6.62e-32) and endothelial cells (OR=4.21; P_FDR_=1.18e-28; Dataset S3).

### MAPT

The strongest association between *MAPT* expression and the atrophy pattern in symptomatic MAPT mutation carriers was measured with microarray probe A_24_P224488, which showed a non-significant positive association z=0.867 (P_FDR_=0.15; Figure 2).

The most significantly negatively correlated gene was *GPR158* (G-protein-coupled receptor 158, Figure 2; Table 2). As in the case of *C9orf72*, the significantly negatively correlated genes showed enrichment for mitochondria (GO_MITOCHONDRIAL_PART; OR=2.11; P_FDR_=2.90e-22) and cellular respiration (GO_CELLULAR_RESPIRATION; OR=3.69; P_FDR_=1.72e-11; Dataset S2). Notably, KEGG pathways for neurodegenerative disorders were enriched (Parkinson’s OR=6.05 P_FDR_=2.75e-16; Alzheimer’s OR=3.37 P_FDR_=7.76e-11; Huntington’s OR=3.08 P_FDR_=4.47e-10). Within the cell types, neuronal genes were strongly enriched (OR=2.26; P_FDR_=2.22e-46).

The most significantly positively associated gene was *LPA* (Lipoprotein(a); Figure 2; Table 2). Significantly positively correlated genes are enriched for genes related to nervous system development (GO_REGULATION_OF_NERVOUS_SYSTEM_DEVELOPMENT; OR=1.83, P_FDR_=3.03e-08) and gliogenesis (GO_GLIOGENESIS; OR=2.66; P_FDR_=2.57e-06). Furthermore, the positively correlated genes were strongly enriched for genes related to mature astrocytes (OR=5.33, P_FDR_=5.66e-102) and moderately enriched for oligodendrocytes related genes (OR=1.78, P_FDR_=2.46e-04; Dataset S3).

Enrichment analyses for GO terms for each genetic group and the consensus list are summarized in Figure 3.

**Figure 3:**
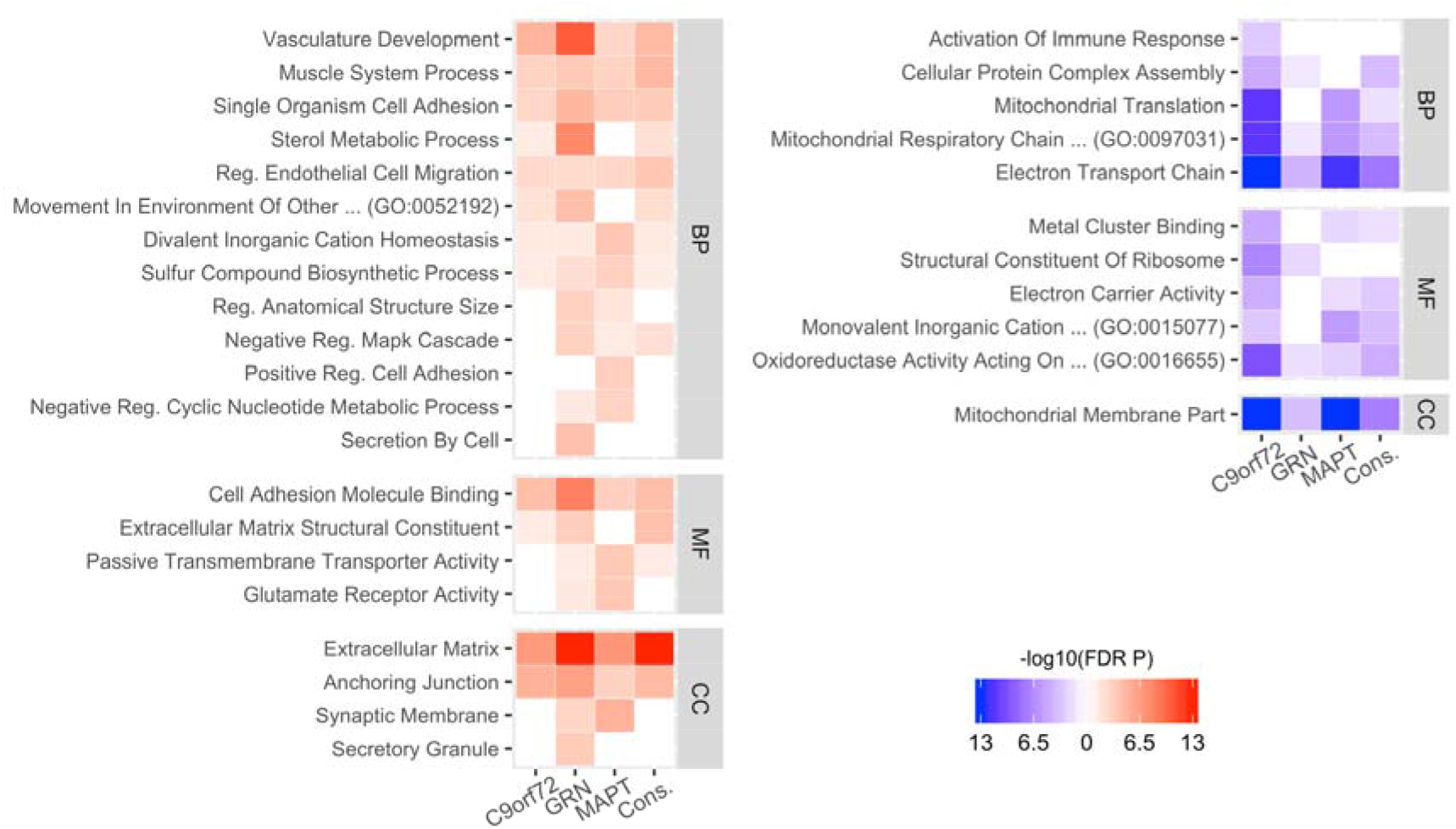
Summary of GO enrichment analysis. Selected GO enrichment results for the three genetic groups and the consensus lists (Cons.). The left panel shows enrichment for genes showing a significant positive correlation with atrophy while the right panel depicts enrichments for negatively correlated genes. Significance of the enrichment is color-coded: more saturated colours signify smaller p-values and non-significant results (i.e., FDR corrected p-value > 0.05) appear white. Depicted GO terms were reduced by filtering based on the number of genes (<500) and by clustering terms with similar meaning using the measure of semantic similarity introduced by Wang *et al*. (2007). The full list of results is available in Dataset S2. BP=biological process, MF=molecular function, CC=cellular component.

### Consensus lists

From the gene lists obtained for each of the FTD gene atrophy maps we created two consensus lists: one comprising 423 genes that were significantly positively correlated with atrophy in all three FTD genes and one list comprising the 700 genes that were significantly negatively correlated in all three maps (Figure 1). Using these lists, we aimed to identify a common theme underlying the atrophy in the three causative genes.

The high-level analysis EWCE showed enrichment for vascular marker genes (Figure 4; Z-score=9.44; P_FDR_=5.99e-05), astrocyte genes (Figure 4; Z-score=3.24; P_FDR_=0.0033) among genes with positive correlation to atrophy severity. Genes negatively correlated with atrophy were highly enriched for neuronal genes (Z-score=6.65; P_FDR_<5.99e-05) and immune cell genes (Z-score=2.56; P_FDR_=0.019). The detailed EWCE analysis of the 189 cell-types confirmed the high-level analysis: nearly all cell-types belonging to the astrocyte and vascular classes showed significant enrichment in the positive consensus list (Dataset S4). The negative consensus list was enriched for microglia and for inhibitory as well as excitatory neurons.

**Figure 4:**
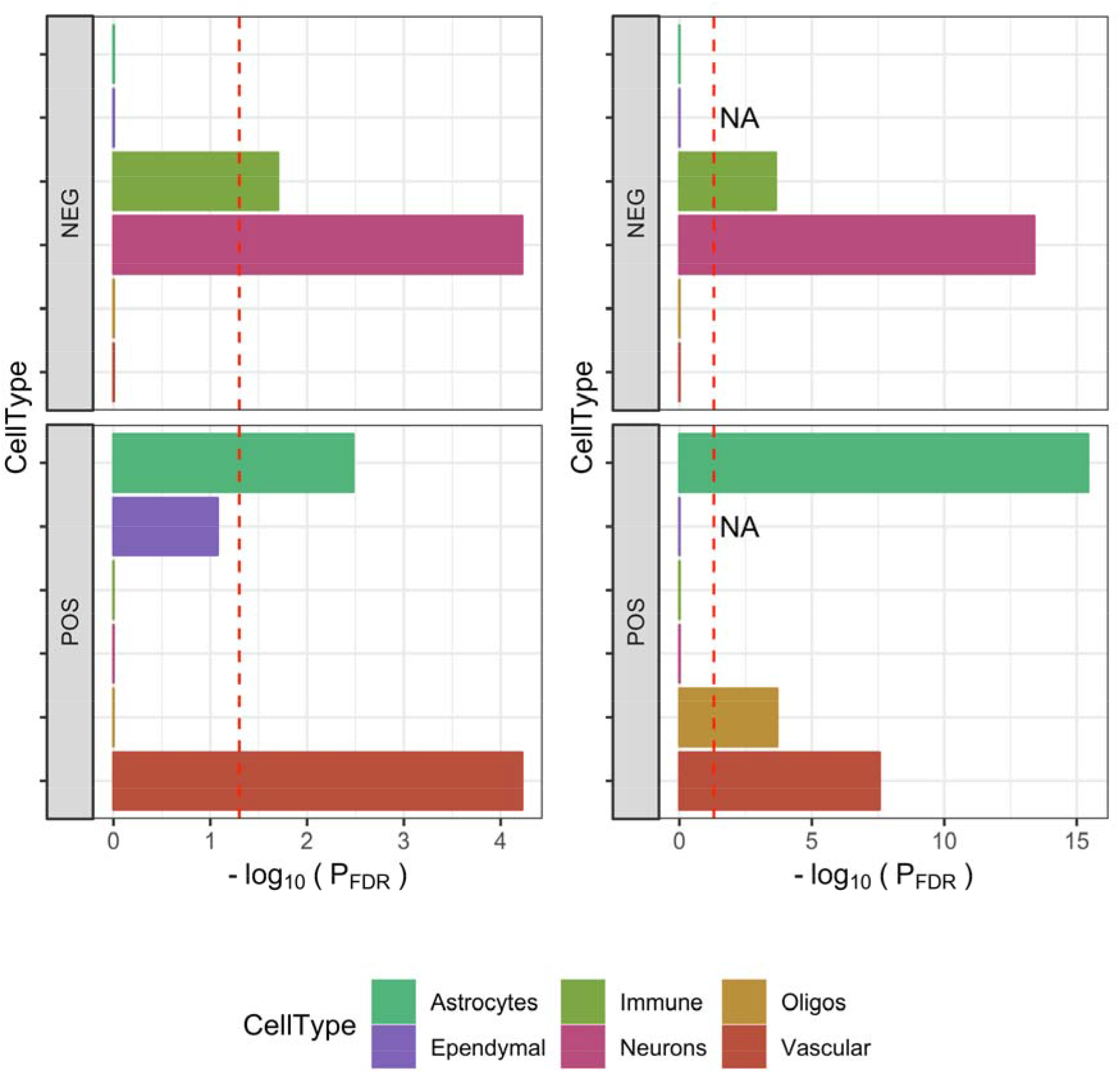
Cell-type enrichment in consensus gene lists. Two different methods were used to investigate cell type signatures in the two consensus lists (POS = gene expression is positively correlated with atrophy; NEG = gene expression is negatively correlated with atrophy). The left panels used the Expression Weighted Cell-type Enrichment (EWCE) method together with single-cell RNA sequencing data from mouse brains (Zeisel *et al*., 2018); the enrichment is summarized as −log_10_(P_FDR_). The right panels used a classic overrepresentation analysis based on Fisher’s exact test with cell-type signature genes obtained from bulk RNA sequencing of purified human cells (Zhang *et al*., 2016). The red dashed lines indicate statistical significance after FDR correction (i.e., −log_10_(0.05)). Both approaches demonstrated a strong neuronal and immune signature in negatively correlated genes (upper panels) and a strong vascular and astrocyte signature in positively correlated genes (lower panels).

These results were further confirmed using cell-type marker genes derived from RNA sequencing of purified human cells (Zhang *et al*., 2016) (Figure 4; right column), with the strongest enrichment being seen for astrocyte marker genes (odds-ratio=3.89; P_FDR_=3.7e-16). Additionally, oligodendrocytes marker genes were enriched among genes with positive correlation to atrophy severity (odds-ratio=2.61; P_FDR_=2.04e-04).

The remaining gene set analyses confirmed results obtained with the individual gene lists (Figure 3; Dataset S2).

## Discussion

We investigated the gene expression correlates of the cortical regions specifically atrophic in the three main genetic causes of FTD. Whilst there was no association with expression of the gene itself (i.e., *C9orf72, GRN* and *MAPT*), our analysis revealed that groups of genes commonly associated with astrocytes and endothelial cells showed higher expression levels in regions with more atrophy and genes commonly associated with neurons and microglia showed higher expression levels in the relatively spared regions.

Astrocytes are the most abundant cell-type in the human CNS and carry out a plethora of functions including biochemical support for the blood-brain barrier-forming endothelial cells, trophic support for neurons, regulation of extracellular ion balance and participation in repair processes of the brain following injuries. Astrocytes reacting to injuries in the central nervous system, reactive astrocytes, are characterized by expression of glial fibrillary acidic protein (GFAP). Depending on the context, such reactive GFAP-positive astrocytes can be neurotoxic (Liddelow and Barres, 2015; Liddelow *et al*., 2017) or neuroprotective (Anderson *et al*., 2016). There are already multiple lines of evidence linking astrocyte (dys-)function to neurodegeneration (Rodríguez *et al*., 2009; Phatnani and Maniatis, 2015; Sofroniew, 2015) and to FTD in particular. For instance, histopathological studies in FTD have shown that severity of astrocytosis and astrocytic apoptosis correlates with the degree of neuronal loss as well as with the stage of the disease (Broe *et al*., 2004). In addition, astrocyte reactivity appears to be region specific in that higher numbers of reactive (GFAP-positive) astrocytes were found in the frontal and temporal cortices of FTD patients compared to controls (Martinac *et al*., 2001). These observations extend to the CSF where levels of GFAP were increased in various neurodegenerative disorders compared to cognitively normal adults with the highest levels in FTD patients (Ishiki *et al*., 2016). *Martinac et al*. (2001) found that degrading astrocytes were inversely correlated with cerebral blood flow in FTD. However, more importantly, astrocytes derived from induced pluripotent stem cells of patients with mutations in *MAPT* were found to demonstrate increased vulnerability to oxidative stress and exhibit disease-associated gene-expression changes (Hallmann *et al*., 2017). Co-culture experiments of such modified FTD astrocytes with previously healthy neurons led, among other things, to increased oxidative stress in these neurons. Hence, taken together, astrocyte reactivity and activation of GFAP may well co-occur with disease onset and progression in FTD.

A second cell-type for which we observed consistent enrichment in brain regions with atrophy were endothelial (or vascular) cells, whilst the overrepresentation analysis was also enriched for vascular terms (circulatory system development). Furthermore, *MXRA8* was found in the top 10 for all three genetic group lists – this gene codes for limitrin, a protein specific to the glia limitans, a component of the blood-brain barrier (BBB). Overall, this suggests a role of the BBB in regional vulnerability in genetic FTD. Dysfunction of the BBB has been implicated in the aetiology of many neurodegenerative disorders (Sweeney *et al*., 2018), with BBB permeability previously found to be abnormal in patients with FTD (Janelidze *et al*., 2017). The BBB is ubiquitous in the cortex, however, our results based on bulk tissue gene expression profiling would suggest regional differences in density of endothelial cells and their link to disease specific atrophy patterns. Indeed, vascular structure in the brain is known to be heterogenous and characterized by differential pathophysiological responses (Cavaglia *et al*., 2001).

Brain regions that were relatively spared in FTD showed higher expression of neuron-related genes. The detailed EWCE analyses showed that both inhibitory and excitatory neuron types were enriched (Dataset S4). Higher expression in spared regions implies lower gene expression in affected regions, which by itself could be a source for increased vulnerability since minor perturbations in gene expression may have a disproportionate effect. However, given that gene expression data was generated using bulk tissue sampling and that neurons have a significant spatial spread via extended dendritic and axonal processes, the observed enrichment signal could originate from differences between mRNA located in the soma and in dendrites, respectively. A recent pathway analysis of 2,028 differentially dendritically localized mRNA isoforms showed strong enrichment for GO terms such as respiratory electron transport chain (Middleton *et al*., 2019), similar to our enrichment pattern. Indeed, a post-hoc enrichment analysis of these dendritically localized mRNA isoforms (Table S2 from (Middleton *et al*., 2019)) confirmed strong enrichment in all three individual gene lists (OR>1.5, P<1.07e-10) as well as the consensus list (OR=1.92, P=5.35e-09). Thus, this result may suggest increased and decreased dendrite density in spared and affected regions, respectively. Neuron morphology is very diverse and notably neurons with selective vulnerability in FTD such as von Economo neurons and fork cells (Kim *et al*., 2012; Seeley *et al*., 2012) are morphologically characterized by large cell bodies and limited branching of dendrites.

In addition to neuron-related genes, there was also an enrichment of immune system-related genes (e.g., microglia genes) in brain regions with reduced atrophy. Recent works using ^⊓^C-PK-11195 PET, a marker of activated microglia, have linked regional patterns of neuroinflammation to protein aggregation in FTD (Bevan-Jones *et al*., 2020; Malpetti *et al*., 2020). Furthermore, a number of genes associated with immune function were strongly negatively correlated with gene expression in the association analysis. In particular, *TREM2* showed the strongest negative correlation in the *C9orf72* list and was also significantly associated in the *GRN* and *MAPT* lists (z< −4.39, P_FDR_<2.68e-04); this is a transmembrane receptor that participates in modulation of the immune response. Heterozygous variants in *TREM2* are known to be a risk factor for Alzheimer’s disease (Guerreiro *et al*., 2013) and levels of the TREM2 protein have been shown to be abnormal in FTD (Heywood *et al*., 2018; Woollacott *et al*., 2018).

Molecular pathways related to mitochondria were also negatively correlated with gene expression in the analysis. Mitochondrial abnormalities have recently been associated with FTD (Lau *et al*., 2018), particularly with *C9orf72* mutations (Lopez-Gonzalez *et al*., 2016; Choi *et al*., 2019), although little is known at present about the exact molecular mechanisms underlying such dysfunction.

The three individual genetic group lists as well as the consensus list feature significant enrichment for KEGG pathways for Alzheimer’s, Parkinson’s and Huntington’s disease. This may be due to a strong overlap in dendritically localized mRNA isoforms and these pathways. Notably, the KEGG pathway for ALS, which features only 53 genes, did not reach statistical significance after FDR correction in any of the three groups (OR=2.12, P=0.0198, P_FDR_=0.59 in the *GRN* list, P>0.05 in the *MAPT* and *C9orf72* lists).

Summarizing, we detected higher levels of astrocyte and endothelial cell-related genes in regions with neurodegeneration in FTD and we found that genes associated with neurons and microglia were more enriched in brain regions that are spared in FTD. Furthermore, there was enrichment for genes that are associated with mitochondria, particularly cellular respiration, in regions that are not affected by atrophy. These results confirm earlier results from an unbiased proteomic screen of tissue samples where the modules related to synapse (M1), mitochondrion (M3) and neuron differentiation (M8) showed negative correlations with clinicopathological traits in FTD, i.e., these three modules were consistently negatively correlated with FTD pathology (Umoh *et al*., 2018). Consistent with our results, there was a positive correlation between clinicopathological traits and modules M5 (Extracellular matrix) and M6 (Response to biotic stimulus), which both showed strong cell-type enrichment for astrocyte-related proteins. Notably, gene sets associated with the extracellular matrix were also enriched in our analysis with regions showing increased atrophy in FTD (Dataset S2). While Umoh *et al*. (2018) interpreted the negative (and positive) correlations in part with a disease-related shift in cell population in the sampled ROIs, our results extend this observation to regional cell type densities since the gene expression samples were obtained from six cognitively normal subjects.

In summary, our analysis of bulk tissue expression data suggests that cortical regions exhibiting the most severe atrophy in genetic FTD may be those with higher expression of astrocyte and endothelial cell-related genes in healthy subjects. Regional cell-density measurements obtained from post mortem histological samples of healthy subjects would be required to confirm this finding. However, our observation fits well with increased blood brain barrier permeability in FTD patients and recent findings of the neurotoxic potential of astrocytes (Liddelow *et al*., 2017). We hypothesize that the distinct regional atrophy pattern in genetic FTD may be driven by regions with naturally increased astrocyte density where these universal astrocyte neurotoxic effects come to bear with higher frequency. Thus, neurodegeneration may be the result of age-related increase in neurotoxic (A1) and senescent astrocytes, which lost many normal astrocytic functions.

## Abbreviations

*C9orf72*: Chromosome 9 open reading frame 72 gene
EWCE: Expression Weighted Cell-type Enrichment
FDR: False Discovery Rate
FTD: Frontotemporal dementia
GENFI: Genetic FTD Initiative
GFAP: glial fibrillary acidic protein
GO: Gene Ontology
*GRN*: Progranulin gene
MAPT: tau gene
MNI: Montreal Neurological Institute and Hospital
OR: Odds Ratio
VBM: Voxel Based Morphometry

## Funding

AA holds a Medical Research Council eMedLab Medical Bioinformatics Career Development Fellowship. This work was supported by the Medical Research Council (grant number MR/L016311/1). The Dementia Research Centre is supported by Alzheimer’s Research UK, Brain Research Trust, and The Wolfson Foundation. This work was supported by the NIHR Queen Square Dementia Biomedical Research Unit, the NIHR UCL/H Biomedical Research Centre and the Leonard Wolfson Experimental Neurology Centre (LWENC) Clinical Research Facility as well as an Alzheimer’s Society grant (AS-PG-16-007). JDR is supported by an Medical Research Council Clinician Scientist Fellowship (MR/M008525/1) and has received funding from the NIHR Rare Disease Translational Research Collaboration (BRC149/NS/MH). This work was also supported by the Medical Research Council UK GENFI grant (MR/M023664/1) and the Bluefield Project. M.R. holds a Medical Research Council Clinician Scientist Fellowship (grant number MR/N008324/1). R.H.R. was supported through the award of a Leonard Wolfson Doctoral Training Fellowship in Neurodegeneration. This work was supported by Italian Ministry of Health (CoEN015 and Ricerca Corrente). Several authors of this publication (JvS, MS, RV, AD, MO, JR) are members of the European Reference Network for Rare Neurological Diseases - Project ID No 739510. This project was supported, in part, via the European Union’s Horizon 2020 research and innovation program grant 779257 “Solve-RD” (to M.S.).

## Supplementary information

**Dataset S1: *Results from the spatial correlation analysis*.** This table shows for every eligible probe, the mapped gene name, the z-score of the meta-analysis for the six donor brains, the corresponding P-values and the FDR adjusted P-value for associations with the three separate t-maps.

**Dataset S2: *Enrichments for pathways*.** Each sheet in this table shows the results of the enrichment analysis for the six gene lists (genes positively and negatively correlated with atrophy, respectively) and the two consensus lists. Columns represent the pathway name, the number of overlapping genes between list and pathway, the expected number of overlapping genes, the odds ratio (OR) and the overrepresentation p-value (including FDR-correction).

**Dataset S3: *Enrichments for brain six cell-types*.** Enrichment analysis testing gene lists with significant positive or negative correlation for cell-type marker genes. Human cell-type gene lists for neurons (N), microglia (MG), mature astrocytes (MA), endothelial cells (EC), oligodendrocytes (OLG) and oligodendrocyte precursor cells (OPC) were obtained from Zhang et al. (2016).

**Dataset S4: *Fine-grained EWCE results* of the two consensus lists**. The columns provide the cell-type abbreviation, the broad cell class (oligodendrocyte, neuron, astrocytes, ependymal, vascular and immune), a more detailed description of the cell type followed by enrichment statistics for the negative consensus and the positive consensus list, respectively. P-values were obtained using 100,000 permutation**s**.

## Appendix

### List of GENFI consortium authors

Caroline Greaves BSc^1^, Georgia Peakman MSc^1^, Rachelle Shafei MRCP^1^, Emily Todd MRes^1^, Martin N. Rossor MD FRCP^1^, Jason D. Warren PhD FRACP^1^, Nick C. Fox MD FRCP^1,2^, Henrik Zetterberg^2^, Rita Guerreiro PhD^3^, Jose Bras PhD^3^, Jennifer Nicholas PhD^4^, Simon Mead PhD^5^, Lize Jiskoot PhD^6^, Lieke Meeter MD^6^, Jessica Panman MSc^6^, Janne M Papma PhD^6^, Rick van Minkelen PhD^7^, Yolanda Pijnenburg PhD^8^, Myriam Barandiaran PhD^9,10^, Begoña Indakoetxea MD^9,10^, Alazne Gabilondo MD^10^, Mikel Tainta MD^10^, Maria de Arriba BSc^10^, Ana Gorostidi PhD^10^, Miren Zulaica BSc^10^, Jorge Villanua MD PhD^11^ Zigor Diaz^12^, Sergi Borrego-Ecija MD^13^, Jaume Olives MSc^13^, Albert Lladó PhD^13^, Mircea Balasa PhD^13^, Anna Antonell PhD^13^, Nuria Bargallo PhD^14^, Enrico Premi MD^15^, Maura Cosseddu MPsych^15^, Stefano Gazzina MD^15^, Alessandro Padovani MD PhD^15^, Roberto Gasparotti MD^16^, Silvana Archetti MBiolSci^17^, Sandra Black MD^19^, Sara Mitchell MD^19^, Ekaterina Rogaeva PhD^20^, Morris Freedman MD^21^, Ron Keren MD^22^, David Tang-Wai MD^23^, Linn Öijerstedt MD^24^, Christin Andersson PhD^25^, Vesna Jelic MD^26^, Hakan Thonberg MD^27^, Andrea Arighi MD^28,29^, Chiara Fenoglio PhD^28,29^, Elio Scarpini MD^28,29^, Giorgio Fumagalli MD^28,29^, Thomas Cope MRCP^30^, Carolyn Timberlake BSc^30^, Timothy Rittman MRCP^30^, Christen Shoesmith MD^31^, Robart Bartha PhD^32,33^, Rosa Rademakers PhD^34^, Carlo Wilke MD^35,36^, Hans-Otto Karnarth MD^37^, Benjamin Bender MD^38^, Rose Bruffaerts MD PhD^39^, Philip Van Damme MD PhD^40^, Mathieu Vandenbulcke MD PhD^41,42^, Catarina B. Ferreira MSc^43^, Gabriel Miltenberger PhD^44^, Carolina Maruta MPsych PhD^45^, Ana Verdelho MD PhD^46^, Sónia Afonso BSc^47^, Ricardo Taipa MD PhD^48^, Paola Caroppo MD PhD^49^, Giuseppe Di Fede MD PhD^49^, Giorgio Giaccone MD^49^, Sara Prioni PsyD^49^, Veronica Redaelli MD^49^, Giacomina Rossi MSc^49^, Pietro Tiraboschi MD^49^, Diana Duro NPsych^50^, Maria Rosario Almeida PhD^50^, Miguel Castelo-Branco MD PhD^50^, Maria João Leitão BSc^51^, Miguel Tabuas-Pereira MD^52^, Beatriz Santiago MD^52^, Serge Gauthier MD^53^, Pedro Rosa-Neto MD PhD^54^, Michele Veldsman PhD^55^, Paul Thompson^56^, Tobias Langheinrich^56^, Catharina Prix MD^57^, Tobias Hoegen MD^57^, Elisabeth Wlasich Mag. rer. nat.^57^, Sandra Loosli MD^57^, Sonja Schonecker MD^57^, Elisa Semler Dr.hum.biol Dipl. Psych^58^, Sarah Anderl-Straub Dr.hum.biol Dipl.Psych^58^, Luisa Benussi PhD^59^, Giuliano Binetti MD^59^, Michela Pievani PhD^59^, Gemma Lombardi MD^60^, Benedetta Nacmias PhD^60^, Camilla Ferrari^60^, Valentina Bessi^60^, Cristina Polito^61^.

### Affiliations

^1^Dementia Research Centre, Department of Neurodegenerative Disease, UCL Queen Square Institute of Neurology, London, UK; ^2^Dementia Research Institute, Department of Neurodegenerative Disease, UCL Institute of Neurology, Queen Square, London, UK; ^3^Center for Neurodegenerative Science, Van Andel Research Institute, Grand Rapids, Michigan, USA.^4^Department of Medical Statistics, London School of Hygiene and Tropical Medicine, London, UK; ^5^MRC Prion Unit, Department of Neurodegenerative Disease, UCL Institute of Neurology, Queen Square, London, UK; ^6^Department of Neurology, Erasmus Medical Centre, Rotterdam, Netherlands; ^7^Department of Clinical Genetics, Erasmus Medical Centre, Rotterdam, Netherlands; ^8^Amsterdam University Medical Centre, Amsterdam VUmc, Amsterdam, Netherlands; ^9^Cognitive Disorders Unit, Department of Neurology, Donostia University Hospital, San Sebastian, Gipuzkoa, Spain; ^10^Neuroscience Area, Biodonostia Health Research Institute, San Sebastian, Gipuzkoa, Spain; ^11^OSATEK, University of Donostia, San Sebastian, Gipuzkoa, Spain; ^12^CITA Alzheimer, San Sebastian, Gipuzkoa, Spain; ^13^Alzheimer’s disease and Other Cognitive Disorders Unit, Neurology Service, Hospital Clinic, Barcelona, Spain; ^14^Imaging Diagnostic Center, Hospital Clínic, Barcelona, Spain; ^15^Centre for Neurodegenerative Disorders, Neurology Unit, Department of Clinical and Experimental Sciences, University of Brescia, Brescia, Italy; ^16^Neuroradiology Unit, University of Brescia, Brescia, Italy; ^17^Biotechnology Laboratory, Department of Diagnostics, Spedali Civili Hospital, Brescia, Italy; ^18^Clinique Interdisciplinaire de Mémoire Département des Sciences Neurologiques Université Laval Québec, Quebec, Canada; ^19^Sunnybrook Health Sciences Centre, Sunnybrook Research Institute, University of Toronto, Toronto, Canada; ^20^Tanz Centre for Research in Neurodegenerative Diseases, University of Toronto, Toronto, Canada; ^21^Baycrest Health Sciences, Rotman Research Institute, University of Toronto, Toronto, Canada; ^22^The University Health Network, Toronto Rehabilitation Institute, Toronto, Canada; ^23^The University Health Network, Krembil Research Institute, Toronto, Canada; ^24^Center for Alzheimer Research, Division of Neurogeriatrics, Department of Neurobiology, Care Sciences and Society, Bioclinicum, Karolinska Institutet, Solna, Sweden; Unit for Hereditary Dementias, Theme Aging, Karolinska University Hospital, Solna, Sweden; ^25^Department of Clinical Neuroscience, Karolinska Institutet, Stockholm, Sweden; ^26^Division of Clinical Geriatrics, Karolinska Institutet, Stockholm, Sweden; ^27^Center for Alzheimer Research, Divison of Neurogeriatrics, Karolinska Institutet, Stockholm, Sweden; ^28^Fondazione IRCCS Ca’ Granda Ospedale Maggiore Policlinico, Neurodegenerative Diseases Unit, Milan, Italy; ^29^University of Milan, Centro Dino Ferrari, Milan, Italy; ^30^Department of Clinical Neurosciences, University of Cambridge, Cambridge, UK; ^31^Department of Clinical Neurological Sciences, University of Western Ontario, London, Ontario Canada; ^32^Department of Medical Biophysics, The University of Western Ontario, London, Ontario, Canada; ^33^Centre for Functional and Metabolic Mapping, Robarts Research Institute, The University of Western Ontario, London, Ontario, Canada; ^34^Department of Neuroscience, Mayo Clinic, Jacksonville, Florida, USA; ^35^Department of Neurodegenerative Diseases, Hertie-Institute for Clinical Brain Research and Center of Neurology, University of Tübingen, Tübingen, Germany; ^36^Center for Neurodegenerative Diseases (DZNE), Tübingen, Germany; ^37^Division of Neuropsychology, Hertie-Institute for Clinical Brain Research and Center of Neurology, University of Tübingen, Tübingen, Germany; ^38^Department of Diagnostic and Interventional Neuroradiology, University of Tübingen, Tübingen, Germany; ^39^Laboratory for Cognitive Neurology, Department of Neurosciences, KU Leuven, Leuven, Belgium; ^40^Neurology Service, University Hospitals Leuven, Belgium, Laboratory for Neurobiology, VIB-KU Leuven Centre for Brain Research, Leuven, Belgium; ^41^Geriatric Psychiatry Service, University Hospitals Leuven, Belgium; ^42^Neuropsychiatry, Department of Neurosciences, KU Leuven, Leuven, Belgium; ^43^Laboratory of Neurosciences, Institute of Molecular Medicine, Faculty of Medicine, University of Lisbon, Lisbon, Portugal; ^44^Faculty of Medicine, University of Lisbon, Lisbon, Portugal; ^45^Laboratory of Language Research, Centro de Estudos Egas Moniz, Faculty of Medicine, University of Lisbon, Lisbon, Portugal; ^46^Department of Neurosciences and Mental Health, Centro Hospitalar Lisboa Norte - Hospital de Santa Maria & Faculty of Medicine, University of Lisbon, Lisbon, Portugal; ^47^Instituto Ciencias Nucleares Aplicadas a Saude, Universidade de Coimbra, Coimbra, Portugal; ^48^Neuropathology Unit and Department of Neurology, Centro Hospitalar do Porto - Hospital de Santo António, Oporto, Portugal; ^49^Fondazione IRCCS Istituto Neurologico Carlo Besta, Milano, Italy; ^50^Faculty of Medicine, University of Coimbra, Coimbra, Portugal; ^51^Centre of Neurosciences and Cell biology, Universidade de Coimbra, Coimbra, Portugal; ^52^Neurology Department, Centro Hospitalar e Universitario de Coimbra, Coimbra, Portugal; ^53^Alzheimer Disease Research Unit, McGill Centre for Studies in Aging, Department of Neurology & Neurosurgery, McGill University, Montreal, Québec, Canada; ^54^Translational Neuroimaging Laboratory, McGill Centre for Studies in Aging, McGill University, Montreal, Québec, Canada; ^55^Nuffield Department of Clinical Neurosciences, Medical Sciences Division, University of Oxford, Oxford, UK; ^56^Division of Neuroscience and Experimental Psychology, Wolfson Molecular Imaging Centre, University of Manchester, Manchester, UK; ^57^Neurologische Klinik, Ludwig-Maximilians-Universität München, Munich, Germany; ^58^Department of Neurology, University of Ulm, Ulm; ^59^Instituto di Ricovero e Cura a Carattere Scientifico Istituto Centro San Giovanni di Dio Fatebenefratelli, Brescia, Italy; ^60^Department of Neuroscience, Psychology, Drug Research, and Child Health, University of Florence, Florence, Italy. ^61^Department of Biomedical, Experimental and Clinical Sciences “Mario Serio”, Nuclear Medicine Unit, University of Florence, Florence, Italy.

## Notes

### Competing Interest Statement

The authors have declared no competing interest.

### Summary of Updates

* revised association analysis * updated discussion * updated figures

## References

Anderson MA, Burda JE, Ren Y, Ao Y, O’Shea TM, Kawaguchi R, et al. Astrocyte scar formation aids central nervous system axon regeneration. Nature 2016; 532: 195–200.

Arloth J, Bader DM, Röh S, Altmann A. Re-Annotator: Annotation Pipeline for Microarray Probe Sequences. PLoS One 2015; 10: e0139516.

Benjamini Y, Hochberg Y. Controlling the False Discovery Rate: A Practical and Powerful Approach to Multiple Testing. J R Stat Soc Ser B 1995; 57: 289–300.

Bevan-Jones WR, Cope TE, Jones PS, Kaalund SS, Passamonti L, Allinson K, et al. Neuroinflammation and protein aggregation co-localize across the frontotemporal dementia spectrum. Brain 2020; 143: 1010–1026.

Broe M, Kril J, Halliday GM. Astrocytic degeneration relates to the severity of disease in frontotemporal dementia. Brain 2004; 127: 2214–2220.

Cardoso MJ, Modat M, Wolz R, Melbourne A, Cash D, Rueckert D, et al. Geodesic Information Flows: Spatially-Variant Graphs and Their Application to Segmentation and Fusion. IEEE Trans Med Imaging 2015; 34: 1976–1988.

Cash DM, Bocchetta M, Thomas DL, Dick KM, van Swieten JC, Borroni B, et al. Patterns of gray matter atrophy in genetic frontotemporal dementia: results from the GENFI study. Neurobiol Aging 2017; 62:191–196.

Cavaglia M, Dombrowski SM, Drazba J, Vasanji A, Bokesch PM, Janigro D. Regional variation in brain capillary density and vascular response to ischemia. Brain Res 2001; 910: 81–93.

Choi SY, Lopez-Gonzalez R, Krishnan G, Phillips HL, Li AN, Seeley WW, et al. C9ORF72-ALS/FTD-associated poly(GR) binds Atp5a1 and compromises mitochondrial function in vivo. Nat Neurosci 2019; 22: 851–862.

Conover WJ, Iman RL. Rank transformations as a bridge between parametric and nonparametric statistics. Am Stat 1981; 35: 124–129.

Dormann CF, McPherson JM, Araújo MB, Bivand R, Bolliger J, Carl G, et al. Methods to account for spatial autocorrelation in the analysis of species distributional data: A review. Ecography (Cop) 2007; 30:609–628.

Grothe MJ, Sepulcre J, Gonzalez-Escamilla G, Jelistratova I, Schöll M, Hansson O, et al. Molecular properties underlying regional vulnerability to Alzheimer’s disease pathology. Brain 2018; 141: 2755–2771.

Guerreiro R, Wojtas A, Bras J, Carrasquillo M, Rogaeva E, Majounie E, et al. TREM2 variants in Alzheimer’s disease. N Engl J Med 2013; 368: 117–127.

Hallmann A-L, Araúzo-Bravo MJ, Mavrommatis L, Ehrlich M, Röpke A, Brockhaus J, et al. Astrocyte pathology in a human neural stem cell model of frontotemporal dementia caused by mutant TAU protein. Sci Rep 2017; 7: 42991.

Hawrylycz MJ, Lein ES, Guillozet-Bongaarts AL, Shen EH, Ng L, Miller JA, et al. An anatomically comprehensive atlas of the adult human brain transcriptome. Nature 2012; 489: 391–399.

Heywood WE, Hallqvist J, Heslegrave AJ, Zetterberg H, Fenoglio C, Scarpini E, et al. CSF proorexin and amyloid-β38 expression in Alzheimer’s disease and frontotemporal dementia. Neurobiol Aging 2018; 72: 171–176.

Ishiki A, Kamada M, Kawamura Y, Terao C, Shimoda F, Tomita N, et al. Glial fibrillar acidic protein in the cerebrospinal fluid of Alzheimer’s disease, dementia with Lewy bodies, and frontotemporal lobar degeneration. J Neurochem 2016; 126: 258–61.

Janelidze S, Hertze J, Nägga K, Nilsson K, Nilsson C, Wennström M, et al. Increased blood-brain barrier permeability is associated with dementia and diabetes but not amyloid pathology or APOE genotype. Neurobiol Aging 2017; 51: 104–112.

Kim EJ, Sidhu M, Gaus SE, Huang EJ, Hof PR, Miller BL, et al. Selective frontoinsular von economo neuron and fork cell loss in early behavioral variant frontotemporal dementia. Cereb Cortex 2012; 22: 251–259.

Lau DHW, Hartopp N, Welsh NJ, Mueller S, Glennon EB, Mórotz GM, et al. Disruption of ER-mitochondria signalling in fronto-temporal dementia and related amyotrophic lateral sclerosis. Cell Death Dis 2018; 9: 327.

Liddelow S, Barres B. SnapShot: Astrocytes in Health and Disease. Cell 2015; 162: 1170.

Liddelow SA, Guttenplan KA, Clarke LE, Bennett FC, Bohlen CJ, Schirmer L, et al. Neurotoxic reactive astrocytes are induced by activated microglia. Nature 2017; 541: 481–487.

Lopez-Gonzalez R, Lu Y, Gendron TF, Karydas A, Tran H, Yang D, et al. Poly(GR) in C9ORF72-Related ALS/FTD Compromises Mitochondrial Function and Increases Oxidative Stress and DNA Damage in iPSC-Derived Motor Neurons. Neuron 2016; 92: 383–391.

Malpetti M, Rittman T, Jones PS, Cope TE, Passamonti L, Jones WRB, et al. In vivo PET imaging of neuroinflammation in familial frontotemporal dementia. medRxiv 2020

Martinac JA, Craft DK, Su JH, Kim RC, Cotman CW. Astrocytes degenerate in frontotemporal dementia: Possible relation to hypoperfusion. Neurobiol Aging 2001; 22: 195–207.

Middleton SA, Eberwine J, Kim J. Comprehensive catalog of dendritically localized mRNA isoforms from sub-cellular sequencing of single mouse neurons. BMC Biol 2019; 17: 1–16.

Modat M, Ridgway GR, Taylor ZA, Lehmann M, Barnes J, Hawkes DJ, et al. Fast free-form deformation using graphics processing units. Comput Methods Programs Biomed 2010; 98: 278–284.

Moran PAP. Notes on Continuous Stochastic Phenomena. Biometrika 1950; 37: 17–23.

Phatnani H, Maniatis T. Astrocytes in neurodegenerative disease. Cold Spring Harb Perspect Biol 2015; 7: a020628.

Richiardi J, Altmann A, Milazzo A-C, Chang C, Chakravarty MM, Banaschewski T, et al. Correlated gene expression supports synchronous activity in brain networks. Sci (New York, NY) 2015; 348: 1241–1244.

Rittman T, Rubinov M, Vértes PE, Patel AX, Ginestet CE, Ghosh BCP, et al. Regional expression of the MAPT gene is associated with loss of hubs in brain networks and cognitive impairment in Parkinson disease and progressive supranuclear palsy. Neurobiol Aging 2016; 48: 153–160.

Rodríguez JJ, Olabarria M, Chvatal A, Verkhratsky A. Astroglia in dementia and Alzheimer’s disease. Cell Death Differ 2009; 16: 378–385.

Rohrer JD, Guerreiro R, Vandrovcova J, Uphill J, Reiman D, Beck J, et al. The heritability and genetics of frontotemporal lobar degeneration. Neurology 2009; 73: 1451–1465.

Seelaar H, Rohrer JD, Pijnenburg YAL, Fox NC, Van Swieten JC. Clinical, genetic and pathological heterogeneity of frontotemporal dementia: A review. J Neurol Neurosurg Psychiatry 2011; 82: 476–86.

Seeley WW, Merkle FT, Gaus SE, Craig AD, Allman JM, Hof PR. Distinctive neurons of the anterior cingulate and frontoinsular cortex: A historical perspective. Cereb Cortex 2012; 22: 245–250.

Skene NG, Grant SGN. Identification of vulnerable cell types in major brain disorders using single cell transcriptomes and expression weighted cell type enrichment. Front Neurosci 2016; 10: 16.

Sofroniew M V. Astrocyte barriers to neurotoxic inflammation. Nat Rev Neurosci 2015; 16: 249–263.

Sweeney MD, Sagare AP, Zlokovic B V. Blood-brain barrier breakdown in Alzheimer disease and other neurodegenerative disorders. Nat Rev Neurol 2018; 14: 133–150.

Umoh ME, Dammer EB, Dai J, Duong DM, Lah JJ, Levey AI, et al. A proteomic network approach across the ALS FTD disease spectrum resolves clinical phenotypes and genetic vulnerability in human brain. EMBO Mol Med 2018; 10: 48–62.

Wang JZ, Du Z, Payattakool R, Yu PS, Chen CF. A new method to measure the semantic similarity of GO terms. Bioinformatics 2007; 23: 1274–1281.

Woollacott IOC, Nicholas JM, Heslegrave A, Heller C, Foiani MS, Dick KM, et al. Cerebrospinal fluid soluble TREM2 levels in frontotemporal dementia differ by genetic and pathological subgroup. Alzheimer’s Res Ther 2018; 10: 79.

Zeisel A, Hochgerner H, Lönnerberg P, Johnsson A, Memic F, van der Zwan J, et al. Molecular Architecture of the Mouse Nervous System. Cell 2018; 174: 999–1014.e22.

Zhang Y, Sloan SA, Clarke LE, Caneda C, Plaza CA, Blumenthal PD, et al. Purification and Characterization of Progenitor and Mature Human Astrocytes Reveals Transcriptional and Functional Differences with Mouse. Neuron 2016; 89: 37.

TECHNICAL WHITE PAPER: MICROARRAY DATA NORMALIZATION [Internet]. 2013[cited 2018 Jul 15] Available from: http://help.brain-map.org/download/attachments/2818165/Normalization_WhitePaper.pdf?version=1&modificationDate=1361836502191&api=v2

